# A Cas12a Toolbox for Rapid and Flexible Group B *Streptococcus* Genomic Editing and CRISPRi

**DOI:** 10.1101/2025.06.20.660720

**Authors:** GH Hillebrand, SC Carlin, EJ Giacobe, HA Stephenson, J Collins, TA Hooven

**Affiliations:** Program in Microbiology and Immunology, University of Pittsburgh School of Medicine, Pittsburgh PA, USA; Department of Pediatrics, Division of Newborn Medicine, University of Pittsburgh School of Medicine, Pittsburgh PA, USA; Dietrich School of Arts and Sciences, University of Pittsburgh, Pittsburgh PA, USA; Department of Microbiology and Immunology, School of Medicine, University of Louisville, Louisville KY, USA; RK Mellon Institute for Pediatric Research, Pittsburgh PA, USA

**Keywords:** Group B *Streptococcus*, genome editing, Cas12a, CRISPRi

## Abstract

*Streptococcus agalactiae* (group B *Streptococcus*; GBS) is a leading cause of neonatal sepsis and meningitis. Despite advances in molecular microbiology, GBS genome engineering remains laborious due to inefficient mutagenesis protocols. Here, we report a versatile and rapid Cas12a-based toolkit for GBS genetic manipulation. We developed two shuttle plasmids—pGBSedit for genome editing and pGBScrispri for inducible CRISPR interference—derived from an *Enterococcus faecium* system and optimized for GBS. Using these tools, we achieved targeted gene insertions, markerless deletions, and efficient, template-free mutagenesis via alternative end-joining repair. Furthermore, a catalytically inactive dCas12a variant enabled inducible gene silencing, with strand-specific targeting effects. The system demonstrated broad applicability across multiple GBS strains and minimal off-target activity, as confirmed by whole-genome sequencing. This Cas12a-based platform offers a rapid, flexible, and scalable approach to genetic studies in GBS, facilitating functional genomics and accelerating pathogenesis research.

## Background

*Streptococcus agalactiae* (group B *Streptococcus*; GBS) is a cause of severe infections, particularly among perinatal populations: pregnant women, newborns, and infants^1,2^. Genetic research is key to the development of GBS vaccines and targeted therapeutics^3–5^.

Currently, most protocols for GBS mutagenesis are based on temperature-dependent shuttle plasmids^6,7^. The process for using a temperature-sensitive mutagenesis plasmid begins with recombinant insertion of an editing cassette with flanking GBS homologous sequences. After transformation into GBS, targeted plasmid integration into the circular chromosome can be promoted by transitioning the culture to a temperature that restricts extrachromosomal plasmid replication^6,8^. In a subsequent self-excision step, the targeted region of the chromosome undergoes a second recombination event in which the native sequence is replaced by the mutagenesis cassette originally introduced on the plasmid^7^.

While temperature-sensitive plasmids have been used to generate many GBS mutants, successful use of these techniques remains time-consuming and laborious. One drawback is that—to propagate the mutagenesis plasmid during the initial cloning and transformant outgrowth steps—the cultures must be maintained at low temperature (usually 28°C), which slows bacterial growth and prolongs the experimental timeframe.

Another problem is that the multiple passaging and outgrowth steps create opportunities for unintended recombination and spontaneous mutation to occur. The result is that, even when using a counterselection-optimized mutagenesis plasmid, it can take weeks or months to create a single targeted GBS mutation. When unexpected problems arise, often the only remedy is to return to the first step of the process, which delays progress.

Mutagenesis approaches based on the activity of clustered regularly interspaced short palindromic repeats and CRISPR-associated protein (CRISPR/Cas) have revolutionized genetic and molecular biology research^9–11^. Multiple CRISPR/Cas system variants evolved in bacteria as defense mechanisms against exogenous, bacteriophage-encoded DNA^12,13^. Bioengineered CRISPR/Cas systems have become useful as easily programmable, targeted molecular effectors of numerous experimental functions including flexible genome editing^14^.

Techniques for using CRISPR/Cas for bacterial genome editing have proven effective^15–18^. However, these efforts can be complicated by the presence of one or more native CRISPR/Cas system encoded on the genome of the study species. In 2023, we introduced a system for using a catalytically inactive dCas9 variant, encoded on the chromosome of bioengineered mutant GBS strains, to downregulate gene expression through CRISPR interference (CRISPRi)^19^. We also made multiple unreported, unsuccessful efforts to use the native GBS CRISPR/Cas system to drive an effective genome editing approach.

An alternative approach to bacterial genome editing is through introduction of non-native bioengineered CRISPR/Cas systems. Cas12a (also referred to as Cpf1) is a type V-A endonuclease found in class 2 CRISPR/Cas systems. Cas12a accepts a smaller guide RNA (gRNA) than Cas9, and unlike the latter, Cas12a gRNA does not require presence of a trans-activating CRISPR RNA (tracrRNA) for programmable targeting of specific DNA sequences^18,20–23^. This allows for smaller, simpler plasmid-based platforms for Cas12a introduction into new experimental systems. Cas12a requires a T-rich protospacer-adjacent motif (PAM) to cleave a target sequence, which makes it an appealing choice for use in low-GC organisms such as GBS^21^.

Recent successful development of a Cas12a-based system for genome editing in *Enterococcus faecalis* prompted us to examine its potential use in GBS^24^. Here we report development, testing, and optimization of a single-plasmid Cas12a-based suite of GBS genetic tools. We have used our Cas12a system for allelic exchange mutagenesis, alternative end-joining mutagenesis, and targeted CRISPRi gene knockdown using a catalytically deactivated Cas12a (dCas12a) variant. Our strategy is efficient. Using it, we have made numerous mutants in an approximately one-week timeframe. Another advantage over other GBS genetic tools (such as our dCas9-based CRISPRi technique) is that it can be used across different GBS strain lineages with no requirement for preceding alterations to the native chromosome. In summary, we find that using Cas12a for site-directed GBS mutagenesis and genetic expression modulation is a significant improvement over previously described techniques.

## Results

### Chromosomal insertion of an eGFP expression cassette using Cas12a and homology-directed repair

Figure 1 diagrams the design and potential uses of the two plasmids that underpin our suite of Cas12a-based GBS genetic tools. pGBSedit is designed for genome editing, while pGBScrispri is for CRISPRi. Both plasmids are modified versions of the Cas12a-based shuttle plasmid pJC005, which was previously developed for *E. faecium* genetic modification^24^. pGBSedit and pGBScrispri both encode conserved gram-negative and gram-positive origins of replication so that CRISPR RNA (crRNA) and homology arm cloning can be performed in *Escherichia coli* followed by electroporation transformation of purified plasmid into GBS.

**Figure 1.**
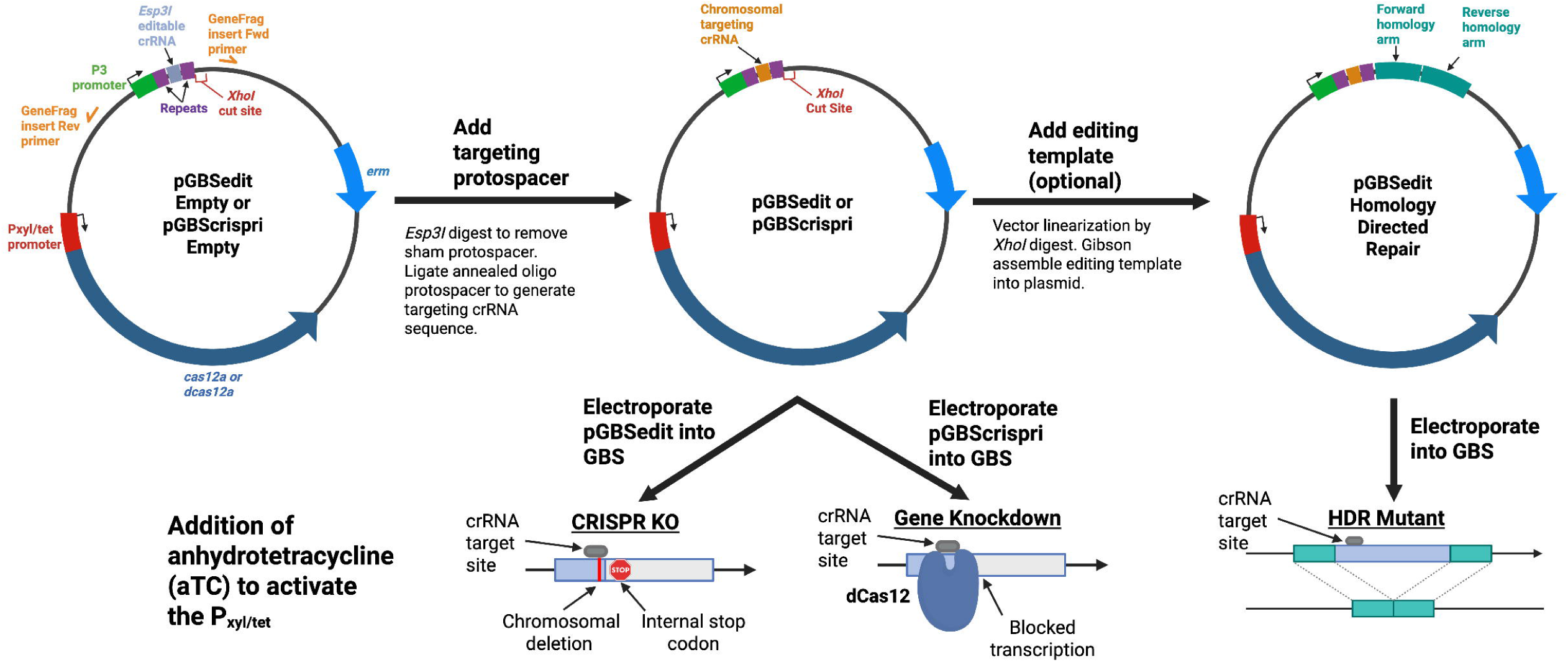
Design and workflow of the Cas12a-based GBS genetic toolkit. Plasmid maps and workflows of the Cas12a-based genetic toolkit for Group B Streptococcus. The two shuttle vectors—pGBSedit (encoding wild-type Cas12a) and pGBScrispri (encoding catalytically inactive dCas12a)—carry a P_xyl/tet_-inducible promoter upstream of the cas12a/dcas12a gene, a constitutive TetR repressor, and an *Esp3I*-flanked protospacer insertion site; pGBSedit also includes a unique *XhoI* site for cloning homology arms. Three experimental protocols are illustrated: (1) homology-directed repair mutagenesis, in which an editing template directs precise genomic insertions or deletions followed by anhydrotetracycline (aTC) selection; (2) template-less mutagenesis, where Cas12a-induced double-strand breaks are repaired via the native alternative end-joining pathway to generate variable indels; and (3) inducible CRISPR interference, in which dCas12a binding to the target locus upon aTC induction leads to targeted gene knockdown.

pGBSedit encodes WT *Acidaminococcus* Cas12a whereas pGBScrispri encodes a modified, catalytically inactive (E993A) dCas12a that can be used for inducible, targeted gene repression. In both versions of the plasmid, the *cas12a*/*dcas12a* gene is downstream of a P_xyl/tet_ inducible promoter bioengineered to maximize the dynamic range in response to its anhydrotetracycline (aTC) inducer, with minimal expression in the uninduced state and strong expression upon aTC exposure^25^. TetR, the promoter’s repressor, is constitutively expressed on the plasmid.

The crRNA cassette is identical in pGBSedit and pGBScripsri. Driven by a constitutive P3 small RNA-optimized promoter, the crRNA region harbors a pair of apposed *Esp3I* restriction enzyme cut sites that, when digested, yield incompatible sticky ends into which a custom dsDNA spacer sequence be ligated to encode a functional crRNA-bearing gRNA.

Once an active crRNA is cloned into place, with genomic target site complementarity adjacent to a Cas12a PAM, aTC induction of WT *cas12a* from pGBSedit leads to reliable selection against all GBS cells bearing the target sequence-PAM combination. The only viable CFU are those that escaped selection by mutation of the target site.

One experimentally useful mechanism of chromosomal repair is homologous recombination with an editing template that can be cloned onto the pGBSedit plasmid at a *XhoI* restriction enzyme recognition site downstream of the crRNA cassette. To demonstrate the effectiveness of this approach, we created an editing template in which the *eGFP* gene and a P23 constitutive promoter is flanked by approximately 500-nt homology arms that direct it to an intergenic region conserved in CNCTC 10/84 and A909 GBS strains.

After cloning this editing template into pGBSedit, along with a targeting crRNA directed to the intended genomic insertion site (generating pGBSedit:eGFP1, Fig. 2A), we transformed WT CNCTC 10/84 and A909 with the targeting plasmid. In the uninduced state without aTC, a liquid culture of the transformed GBS strain grew as an unselected lawn on solid agar media (**Fig. 2B-C, left side**). However, when exposed to aTC added atop the agar media, there is dramatic and reliable selection against the WT background that comprises most of the lawn (**Fig. 2B-C, right side**). The only colonies that grew from the induced CNCTC A909+pGBSedit:eGFP1 and 10/84+pGBSedit:eGFP1 populations expressed eGFP genotypically (Fig. 2D-E) and phenotypically (Fig. 2F-G).

**Figure 2.**
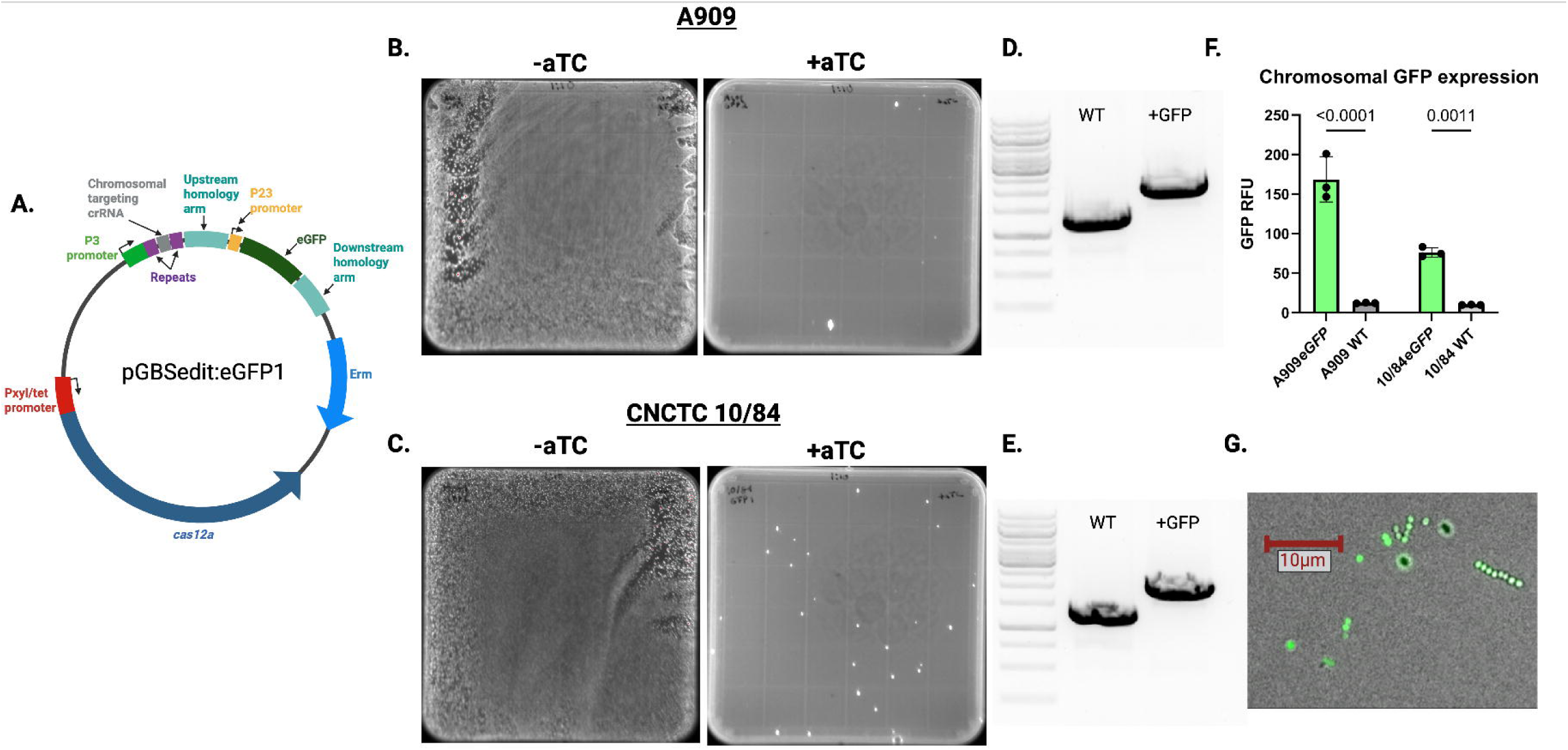
Cas12a-mediated insertion of eGFP into the GBS chromosome via homology-directed repair. (A) Diagram of pGBSedit:eGFP1, with ∼500-nt upstream and downstream homology arms flanking the *eGFP* coding sequence targeted to an intergenic locus conserved in A909 and CNCTC 10/84. (B & C) Representative agar plates of A909 (B) and CNCTC 10/84 (C) transformants on erythromycin: without aTC (left) showing unselected background, and with 500 ng/mL aTC (right) showing only *eGFP* integrants. (D & E) Colony PCR of aTC-selected colonies: expected band sizes for WT (upper) and eGFP insertion (lower) indicated. (F & G) Fluorescence imaging under blue-light illumination confirming *eGFP* expression in selected A909 (G) and CNCTC 10/84 resuspended colonies.

### Markerless gene deletion using Cas12a homology-directed repair

An anticipated use of the Cas12a tools in GBS is markerless gene (or other genomic region) deletion. To demonstrate feasibility of this application and to quantify various possible Cas12a selection endpoints, we created pGBSedit:Δ*covR1* to create a markerless deletion of the *covR* gene. CovR is part of the two-component signaling system that regulates numerous GBS genes^26,27^ , including the *cyl* operon responsible for the biosynthetic pathway of β-hemolysin/cytolysin (βHC), a pigmented, hemolytic GBS toxin^28,29^. We chose this target gene because its deletion leads to *cyl* de-repression and an easily assayed hyperpigmented, hyperhemolytic phenotype^30^.

pGBSedit:Δ*covR1* has an anti-*covR* targeting crRNA directed to the target sequence next to the 3’-TTTC PAM at nucleotide position 436 in the *covR* gene. In the pGBSedit *XhoI* site, we cloned an approximately 1000-bp editing template composed of fused *CovR* upstream and downstream homology arms (Fig. 3A).

**Figure 3.**
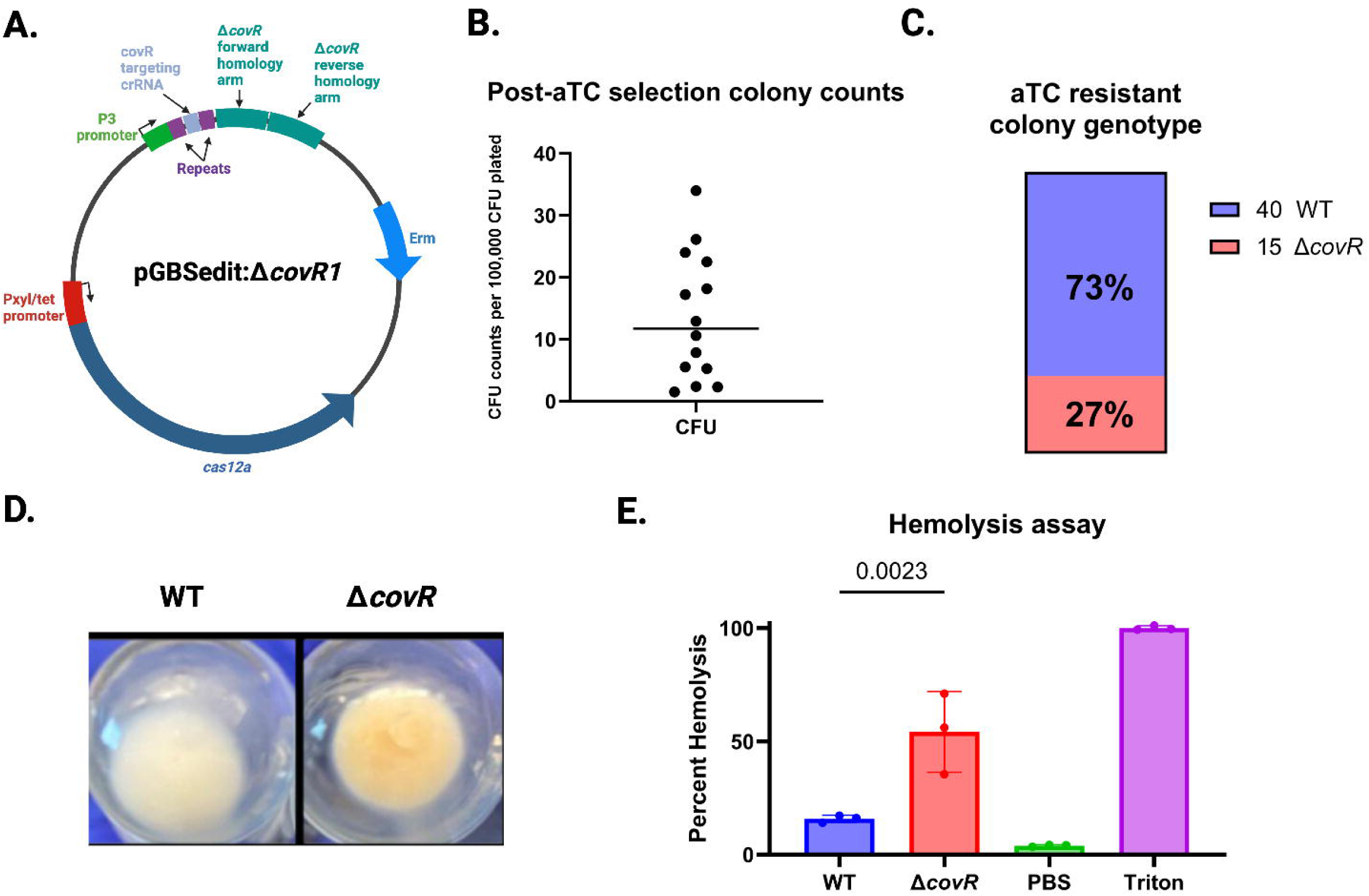
Markerless deletion of covR using Cas12a homology-directed repair. (A) Design of pGBSedit:ΔcovR1, containing fused ∼1 kb homology arms upstream and downstream of the *covR* coding sequence. (B) Selection escape frequency for ΔcovR1 across 14 independent aTC selections in A909; each symbol represents one biological replicate, and the horizontal bar denotes the mean (1.1 × 10⁻⁴). (C) Genotyping by colony PCR of 55 aTC-selected colonies from six biological replicates; 27% yielded the expected ΔcovR amplicon. (D) Photograph of liquid cultures showing hyperpigmented phenotype of confirmed ΔcovR mutants. (E) Quantification of hemolytic activity for Δ*covR* versus WT, measured by released hemoglobin absorbance (n = 3; mean ± SD).

After transformation of purified pGBSedit:Δ*covR1* into WT A909, we performed 14 independent aTC selection experiments during which we quantified the unselected background CFU concentration and the aTC selection output. The mean rate of selection escape across our 14 biological replicates was 1.1×10^-4^ (range 1×10^-5^ – 3.5×10^-4^, Fig. 3B). We performed colony PCR on colonies from 8 independent biological replicates of the transformation step. Among the 55 PCR reactions that generated an electrophoresis gel band, 15 (27%) were of the expected size for the in-frame gene deletion, while the remaining were consistent with wild type (Fig. 3C). Inspection of the 15 suspected knockouts showed that all had the expected hyperpigmented appearance when grown in liquid culture (Fig. 3D). To confirm that the knockout strain was hyperhemolytic, we performed three biological replicates of hemolysis assays with one of the genotypic knockouts from our experiment, which confirmed the expected phenotype (Fig. 3E).

### CRISPR/Cas12a GBS mutagenesis by recombinant free end repair

To examine whether Cas12a-mediated mutations at the genomic target site occur in the absence of homologous recombination with an editing cassette, we generated *covR* and *eGFP* targeting versions of pGBSedit without homology-based editing templates. pGBSedit:*covR2* and pGBSedit:*eGFP2* target the *covR* and *eGFP* coding sequences, respectively; pGBSedit:P23 targets the P23 promoter upstream of *eGFP* (**Fig. 4A-B**). We transformed WT A909 with pGBSedit:*covR2* and A909 with chromosomal eGFP (A909eGFP) with pGBSedit:*eGFP2* or pGBSedit:P23. After selection with aTC, we examined the surviving colonies for phenotypic changes, noting that the majority of the WT A909+pGBSedit:*covR2* colonies were hyperpigmented and hyperhemolytic and the A909eGFP+pGBSedit:*eGFP2* and A909eGFP+pGBSedit:P23 colonies were not fluorescent. We performed colony PCR of the genomic regions flanking the three Cas12a target sites, then Sanger sequenced the resulting bands. This analysis identified two genomic deletions surrounding the Cas12a targeted sites in the *covR* targeted colonies (Fig. 4C). One deletion was 2 bp in size and the other was 60 bp. Sanger sequencing revealed one genomic deletion each for the P23 promoter and *eGFP* targeted strains (**Figs. 4D**). Predictably, mutations that delete significant portions of the P23 promoter and coding sequence frameshift mutations lead to loss of gene function in phenotypic testing (**Fig 4E-F**).

**Figure 4.**
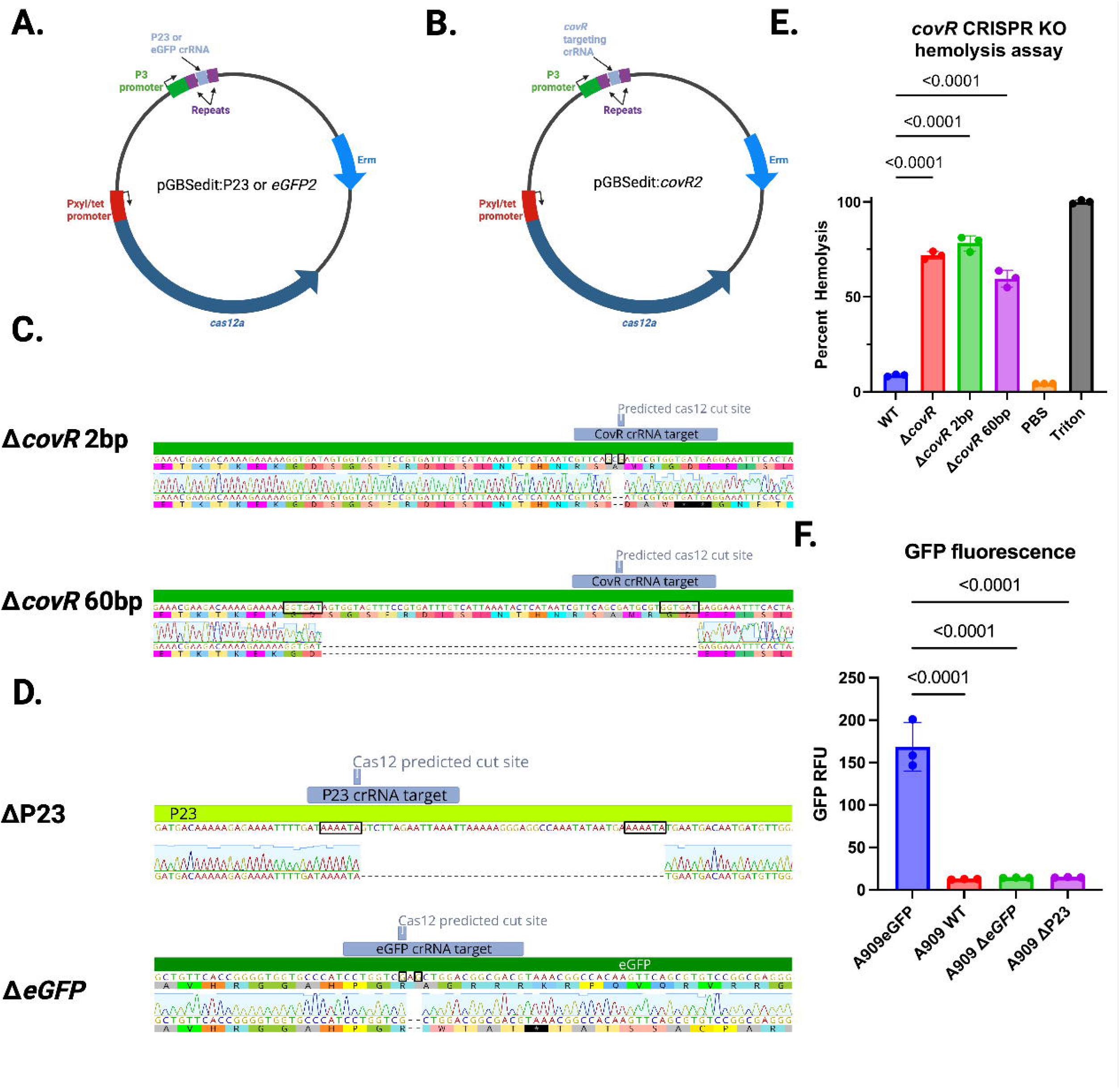
Figure 4. CRISPR/Cas12a-mediated free end-joining mutagenesis in GBS. (A & B) Plasmid designs for template-less mutagenesis: pGBSedit:covR2, pGBSedit:eGFP2, and pGBSedit:P23, each encoding a protospacer targeting covR, chromosomal eGFP, or the P23 promoter. (C) Representative sequence alignments from covR2 mutants showing a 2-bp deletion and a 60-bp deletion at the target site. Microhomology regions suspected to have served as loci for alternative end-joining are boxed. (D) Indel patterns at the P23 and eGFP protospacer sites in selected mutants. Microhomology regions suspected to have served as loci for alternative end-joining are boxed. (E) Hyperhemolytic phenotype of covR mutants in a hemolysis assay (n=3; mean ± SD). (F) Loss of eGFP fluorescence in eGFP2 mutants and reduced P23 promoter activity in P23 mutants.

While the mutations that emerge from this template-less CRISPR editing approach are unpredictably sized, we find that the speed and ease with which they can be generated make this an attractive protocol for rapid creation of mutants in regions of interest. In our benchmarking experiments, the full pipeline of crRNA cloning, GBS transformation and aTC selection, and PCR/Sanger sequence confirmation could all be completed in a week. Promising genomic changes made through this approach can then be more rigorously verified by markerless deletion or allelic exchange with an editing template.

### Whole-genome sequencing of Cas12a-edited GBS genomes does not indicate substantial off-target activity

A persistent concern with CRISPR-based genomic editing is the potential for off-target mutagenesis. Cas12 editing systems has been shown to have promiscuous nickase activity^31^ that could drive higher than baseline rates of mutation if erroneous repair mechanisms were widely activated. To investigate this possibility, we performed Illumina whole-genome sequencing of mutant strains recovered and tested in the above experiments. After unbiased *de novo* alignment and contig scaffolding against a reference GBS genome, we identified between 2 and 9 SNPs across our sequenced mutant strain genomes. Additionally, a prophage resident in the A909 genome was observed to have excised in one of our sequenced strains, which we have encountered in previous studies unrelated to Cas12a mutagenesis^19^. Given estimated secular mutation rates in GBS of approximately 10^-9^ mutations per bp per generation^32–35^, or approximately 0.002 mutations per 2M-bp genome per generation, and roughly 20 generations per laboratory outgrowth, 2-9 fixed SNPs in a strain that is 5-10 outgrowth steps from its progenitor roughly matches expectations, and does not indicate widespread or biased off-site mutagenesis activity by Cas12a.

### Inducible gene silencing with dCas12a

To explore the potential of using dCas12a as a tool for inducible gene silencing, we used crRNA targeting βHC biosynthetic enzymes encoded by the *cyl* operon and crRNA targeting *covR* (Fig. 5A). Previous studies of dCas12a-based gene silencing have demonstrated strand bias in which crRNA targeting the template strand of the gene have a stronger silencing effect than crRNA targeting the non-template strand^36^. This is the opposite of dCas9 gene silencing, which works best when the non-template strand is targeted by the guide RNA (dCas9 template strand targeting can actually increase gene expression in some cases)^19,37^. To examine this aspect in our system, we used both template and non-template targeting crRNA.

**Figure 5.**
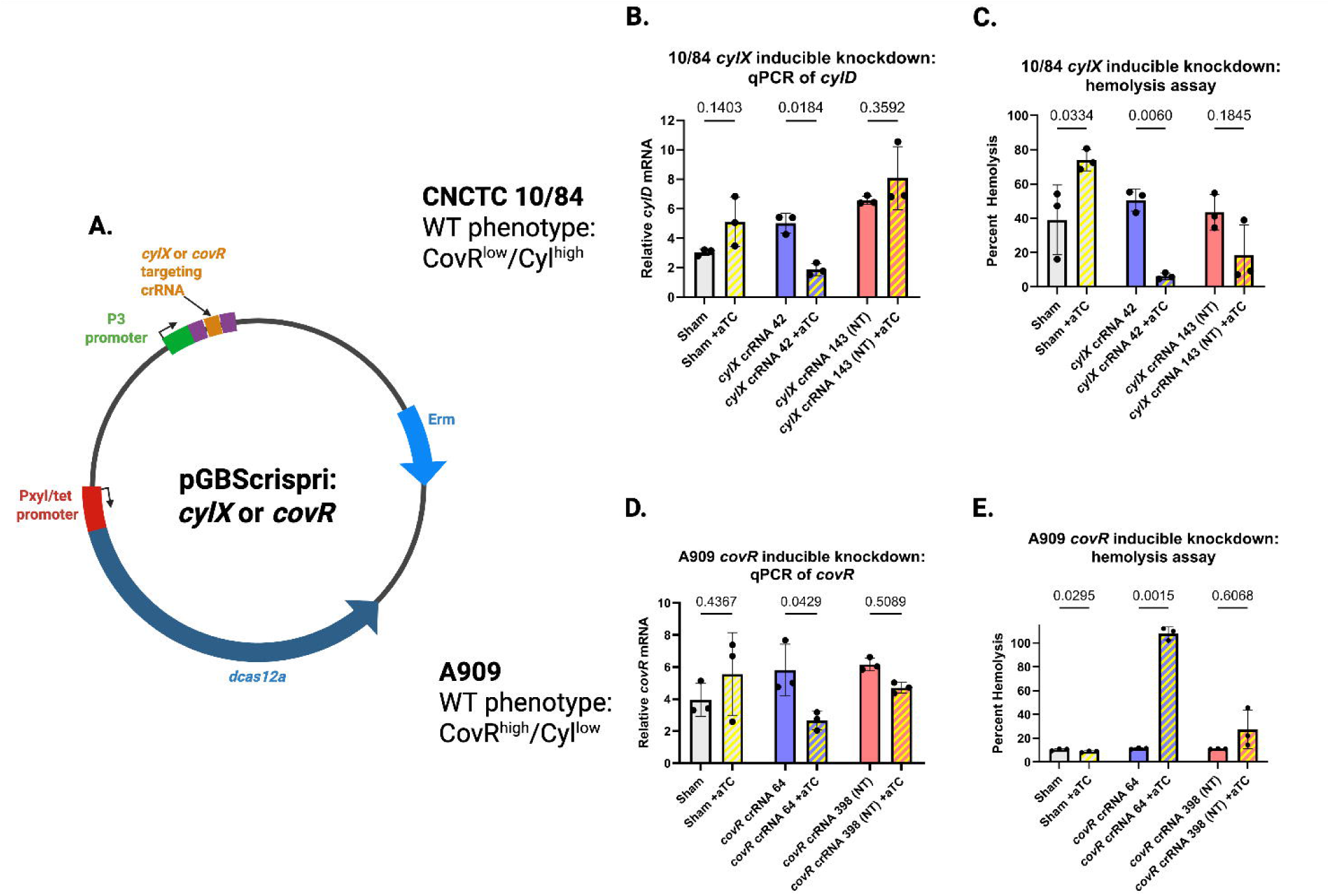
Inducible gene silencing in GBS using dCas12a CRISPR interference. (A) Protospacer designs for dCas12a targeting: cylx42 (template strand) and cylx143 (non-template strand) in the *cyl* operon, and a *covR*-targeting protospacer. (B) qRT-PCR quantification of *cyl* expression in CNCTC 10/84 with ( + aTC) or without ( – aTC) induction (n = 3; mean ± SD), showing stronger knockdown with template-strand targeting. (C) Hemolysis assays of CNCTC 10/84 strains demonstrating reduced βHC activity upon dCas12a targeting of *cyl* (mean ± SD; n = 3). (D) qRT-PCR of *cyl* expression in A909 harboring *covR*-targeting dCas12a, showing de-repression upon induction (n = 3; mean ± SD). (E) Hemolysis assays of A909 *covR* knockdown strain confirming increased βHC activity with aTC induction (mean ± SD; n = 3).

We transformed CNCTC 10/84 with pGBScrispri:*cylx42* (template targeting) and pGBScrispri:*cylx143* (non-template targeting). CNCTC 10/84 is a naturally hyperpigmented/hyperhemolytic GBS strain, due to a SNP in the *covR* promoter that results in constitutively low expression^38^, so suppressing *cyl* in this background is expected to reduce pigmentation and cytotoxicity. By qRT-PCR (Fig. 5B) and *in vitro* hemolysis assays (Fig. 5C), we observed *cyl* downregulation in the presence of the aTC inducer and a targeting crRNA. Addition of aTC without a targeting protospacer was noted to cause a small but consistent upregulation of *cyl* expression, both by qRT-PCR and phenotypic assay. As previously described, the template strand targeting pGBScrispri:*cylx42* plasmid led to greater gene repression than the non-template targeting pGBScrispri:*cylx143* variant.

Silencing *covR* in A909, which is not a naturally hyperhemolytic strain, is expected to cause *cyl* de-repression (Fig. 5D) and overexpression of the βHC toxin. Again, this expected result was observed by qRT-PCR (Fig. 5E) and hemolysis assays (Fig. 5F), only in the presence of the aTC inducer. (In contrast to CNCTC 10/84, aTC did not measurably increase βHC expression in A909 when no *covR* targeting crRNA present, see Fig. 5F leftmost two columns). Together, these results demonstrate the utility of inducible dCas12a as a rapid and reliable tool for targeted gene expression knockdown in GBS.

## Discussion

We have described a suite of Cas12a-based molecular tools to facilitate genetic experiments in GBS. While Cas9-based platforms have dominated the first decade of widespread CRISPR/Cas mediated biology, other Cas variants have been found to be superior in some circumstances. The type V-A Cas12a enzyme, variants of which were first characterized in *Acidaminococcus* spp. and *Lachnospiraceae* spp., generates staggered DNA cleavage ends 18 nt (on the crRNA homologous strand) and 23 nt (on the crRNA complementary strand) from the PAM^21^. This is different from Cas9, which generates blunt-end double-stranded DNA cleavage^12^. Also distinct from Cas9 are the location and sequence requirements of the Cas12a PAM. Whereas Cas9 uses a 5’ PAM, which in most *Streptococci* has a 5’-NGG sequence requirement, Cas12a recognizes 3’-TTTV upstream of the target sequence.

Two major advantages of Cas12a-based technology for altering the GBS genome or gene expression are that 1) the necessary Cas12a molecular components can be encoded on a smaller plasmid than Cas9-based systems, making the cloning and GBS transformation steps more likely to succeed; 2) because of significant structural differences between Cas12a and Cas9, the native GBS CRISPR/Cas9 system has no crosstalk with exogenously introduced Cas12a molecules. Leveraging these two important advantages has allowed us to develop a set of efficient, inexpensive tools for rapid GBS genetic experimentation. The resources described here should allow investigators to choose from several complementary approaches for GBS chromosomal manipulation.

The dCas12a-based CRISPRi system is effective for inducible targeted gene knockdown, as demonstrated by our successful downregulation of the *cyl* operon and the *covR* gene that encodes a key transcriptional regulator of *cyl* and its expression of the hemolytic toxin β/HC. In experiments with our recently published dCas9-based CRISPRi system, we have shown that CRISPRi efficacy is variable^19^. Targeting dCas9 to the 5’ end of a protein-coding gene generally has a greater knockdown effect than targeting closer to the 3’ gene terminus, but even different protospacers directing dCas9 to the first half of a protein coding sequence can cause variable decreases in gene expression. We expect similar findings with dCas12a-based CRISPRi and recommend that—whenever possible— researchers examining gene function with dCas12a employ multiple different protospacers targeting their gene of interest to boost the odds of maximizing the knockdown effect with one of them.

The capabilities around direct chromosomal mutagenesis afforded by our system open new possibilities in terms of speed and ease of generating new genetic variants. The protocol for using a crRNA-only plasmid construct to generate variable end-joining mutations at the target site can be completed in a week, which is a significant advance over temperature-based plasmid selection/counter-selection systems. The recombinant DNA mechanism underlying this end-joining mutagenesis phenomenon is not fully clear.

CRISPR/Cas-based genome editing in eukaryotic cells can make use of non-homologous end-joining (NHEJ) mechanisms, but the enzymatic systems responsible for NHEJ are not present in bacteria. Mechanistically simpler NHEJ processes, which depend on ATP-dependent ligase-D and Ku enzymes, exist in *Mycobacteria* and *Bacillus* species, but are not encoded by GBS^39–41^.

A different bacterial double-stranded break repair pathway, known as alternative end-joining (A-EJ) appears more likely to be functioning in the repair-driven mutagenesis we observe in GBS following chromosomal Cas12a targeting^42,43^. A-EJ is a process of microhomology-based DNA recombination. First described in *E. coli*, A-EJ of bacterial chromosomal breaks is partially dependent on the RecBCD complex driving free end degradation, leading eventually to exposure of compatible microhomologous sequences (approximately 3-nt on average) that drive initial re-synapsis of apposing ends, followed by gap repair. Because of the initial bidirectional free end degradation that underpins A-EJ, the mutations left behind in the process reveal short or long sequence deletions eventually resolved by stochastic microhomology-driven reannealing of the two free ends. Small single-stranded gaps left over after microhomology synapsis are suspected to be repaired by the essential enzyme LigA, followed by resolution of ssDNA flaps by 3’ exonucleases or SbcCD^44^. Because targeted Cas12a-initiated A-EJ creates variable, often frameshifted chromosomal deletions, it can be harnessed as a tool for bioengineering experimental mutations in GBS as we have shown here. A predicted mechanism for GBS A-EJ is diagrammed in **Supplemental Fig. S1**.

Cas12a selection against WT chromosomal sequences can also be used to screen for allelic exchange events generated by homology-driven recombination with an editing template cloned into the same pGBSedit plasmid that carries the Cas12a and gRNA coding sequences. This approach mirrors widely used CRISPR/Cas genome editing strategies that have been effectively harnessed for studies on organisms across the tree of life. We have successfully used Cas12a allelic exchange mutagenesis to replace genes with antibiotic resistance markers, generate markerless gene deletions and SNPs, and insert new sequences at specific sites on the chromosome.

This work highlights the value of flexible tool design when working in genetically diverse species like GBS. The pGBSedit and pGBScrispri plasmids were constructed with modular components that facilitate rapid adaptation to different experimental aims, whether targeting chromosomal loci for allelic exchange or downregulating gene expression through inducible repression. Because these tools do not rely on prior genomic alterations, they are well-suited to applications across strain backgrounds and experimental conditions.

Although this system streamlines many aspects of mutagenesis and gene regulation in GBS, certain limitations remain. As with other CRISPR-based systems, off-target effects cannot be entirely ruled out. Our tetracycline-inducible Cas12a system allows suppression of the enzyme until a temporary exposure drives selection against the target (wild type) genetic sequence, limiting the duration of susceptibility to Cas12a off-target effects.

Accordingly, our whole-genome sequencing results for the mutants we generated in this study did not suggest indiscriminate mutagenesis from widespread Cas12a nickase activity. However, the possibility of unintended mutations remains; whole-genome sequencing or complementary genetic controls are therefore advisable when feasible. In addition, while the ease of using end-joining mutagenesis is appealing, its reliance on stochastic microhomology limits its precision, and it may not be appropriate for applications requiring tightly defined genomic edits.

In summary, the Cas12a-based genetic toolkit presented here provides a set of versatile, efficient alternatives to more cumbersome traditional methods for GBS manipulation. By supporting targeted mutagenesis, promoter knockdown, and recombination-driven genome editing, these tools may facilitate broader efforts to interrogate GBS gene function and pathogenesis. We anticipate that their continued refinement and use will help advance genetic research in this clinically important organism.

## Methods

### Ethics statement

Whole blood collection by phlebotomy of healthy adult volunteers was conducted in accordance with an approved University of Pittsburgh IRB protocol (CR19110106-001). Following use of blood in GBS hemolysis assays, samples were discarded and no further volunteer-level data were collected.

### Reagents

LB liquid broth mix, LB agar, TS liquid broth mix, TS agar, erythromycin, anhydrotetracycline, M17 liquid medium mix, glycine, *Esp3I*, T4 ligase, T4 polynucleotide kinase, *XhoI*, Gibson assembly master mix, *DpnI*, Q5 Taq polymerase, Qiagen QIAprep Spin Miniprep Kit, M17 broth, PEG6000

### Bacterial strains and growth conditions

GBS strains A909 (serotype Ia, sequence type 7) and CNCTC 10/84 (serotype V, sequence type 26), both of which were originally isolated from infected neonates^38,45,46^, and their derivatives were grown at 37°C under stationary conditions in tryptic soy (TS) medium (Fisher Scientific cat. # DF0370-17-d). GBS was supplemented with 5 μg/mL erythromycin as needed for selection. *E. coli* strain DH5α was grown at 37°C with shaking in Luria-Bertani (LB) medium (Fisher Scientific cat. # DF9446-07-5) supplemented with 300 μg/mL erythromycin as needed for selection.

### Statistical analyses

Statistical analyses were performed with GraphPad Prism v. 10.2.3 for macOS.

### Construction of pGBSedit and pGBScrispi

Plasmid backbone pJC005 was used as a backbone for pGBSedit and pGBScrispri. A modular protospacer expression cassette with a small RNA promoter (P3) driving the expression of the repeats and *Esp3I* editable sham protospacer was added to both vectors. Digestion with *Esp3I* allows for complete removal of the sham protospacer and ligation of new protospacers as annealed DNA oligos with matching sticky ends. A unique *XhoI* cut site was also added at the end of this cassette for linearization of the plasmid in use with Gibson Assembly addition of homology arms. This modular cloning cassette was designed and ordered as a gene fragment from Twist Bioscience added to both vectors via Gibson assembly.

### Protospacer design and insertion

Cas12a-compatible GBS genomic target sites with TTTV PAM sequences were identified with the aid of the CRISPick server hosted by the Broad Institute^47^. Under the “CRISPRko/AsCas12a” settings, CRISPick generates a ranked list of 23-nt sgRNA candidate sequences that interface correctly with our GBS Cas12a tools. For dCas12a CRISPRi, sgRNA sequences complementary to the plus-strand should be selected; for Cas12a mutagenesis, plus- or minus-strand sgRNA sequences are equally effective.

After selecting a sgRNA target, the 23-nt sequence can be used to design custom ssDNA protospacer sequences that can be annealed into a dsDNA cassette for cloning into digested pGBSedit or pGBScrispri. The forward oligo is constructed as [TAGAT + sgRNA + A] and the reverse oligo is constructed as [AAATT + reverse complement sgRNA + A]. These ssDNA oligo constructs can be ordered separately from a synthetic DNA provider.

Purified pGBSedit (or pGBScrispri) plasmid, prepared from *E. coli* liquid culture is digested with *Esp3I* according to manufacturer instructions, then the linearized plasmid is gel extracted from an agarose electrophoresis gel.

The forward and reverse protospacer oligonucleotide samples are end-phosphorylated then annealed as follows: rehydration of lyophilized samples to 100 μM stocks then admixture of 1 μL forward oligo, 1 μL reverse oligo, 2.5 μL 10x T4 ligase buffer, 0.5 μL T4 Polynucleotide kinase, 20 μL ultrapure H_2_O. Incubate at 37°C for 1 hour. After this incubation, add 2.5 μL sterile 1 M NaCl, then heat the mixture to 95 °C in a heat block for 5 minutes. After this denaturation step, remove the heat block and allow it to cool slowly to 4°C with the oligo mixture tube remaining inserted in place.

Once phosphorylated and annealed, the dsDNA protospacer can be ligated into *Esp3I*-digested, gel-purified pGBSedit/crispr plasmid in a 20 μL reaction containing 5 μL *Esp3I*-digested plasmid; 2 μL phosphorylated/annealed protospacer; 2 μL 10x T4 ligase buffer, 1 μL T4 ligase, 9 μL H_2_O. Allow this reaction mixture to incubate at 16 °C for at least two hours (overnight incubation is acceptable).

We use 5 μL of this ligation reaction to transform chemically competent *E. coli*. Colony screening by PCR or sequencing-based methods can be used to confirm appropriate protospacer insertion. We perform colony PCR using the forward ssDNA oligo and pGBSedit_crRNA_Screen_R (GCTATCTTCGTCATAGTTACCT).

### Editing template design and insertion

Editing templates included homology arms with approximately 500 bp of overlap (on each end of the intended deletion or insertion). The Cas12a target site, determined with the aid of CRISPick, was positioned roughly equidistant from the homology arms. Editing constructs were ordered as synthesized gene fragments but could alternatively be generated through recombinant techniques depending on the intended end-use. We cloned the editing construct into linearized *XhoI*-digested pGBSedit using Gibson assembly and used PCR and Sanger or whole-plasmid sequencing techniques to verify the insert.

An alternative approach is to synthesize a gene fragment that contains the sgRNA protospacer sequence and an editing template. Such a fragment can be quickly and fully customized. For cloning of a gene fragment into pGBSedit, we use GeneFrag Insert Fwd (GCAATAAAACTTATTCGGATCCGTCGTTTTACAACGTC) and GeneFrag Insert Rev (AGCTGACGGTCAGAGAGAAAGGATTTCTCACATAAAATAGAGG) to linearize pGBSedit using Q5 Taq polymerase. The PCR product can then be treated with *DpnI* to digest circular plasmid, followed by gel or column purification of the PCR reaction mixture. We designed custom gene fragments with homology arms that enable Gibson assembly insertion into linearized pGBSedit, then confirmed proper cloning by whole-plasmid sequencing.

### Plasmid purification and transformation into GBS

Once cloning products had been sequence-confirmed in *E. coli*, plasmids were purified using the Qiagen QIAprep Spin Miniprep Kit according to manufacturer instructions.

Electrocompetent GBS was prepared by seeding a 30 mL culture of M17 broth+ 0.5% glucose (GM17) with a single colony from an overnight agar plate and allowing overnight growth at 37 °C. The next morning, the overnight culture was added to 200 mL of GM17 supplemented with 2% glycine and pre-warmed to 37 °C. The culture growing in glycine-containing broth was monitored until it reached an O.D._600_=0.6. The culture was then pelleted and washed twice in ice-cold 10% polyethylene glycol (average molecular weight=6000; PEG6000), then washed again and the entire pellet resuspended in 25 mL of ice-cold PEG6000 wash solution containing supplemental 25% glycerol. 50 μL aliquots of this suspension were then used for electroporation transformation of the purified shuttle plasmid, followed by overnight outgrowth on selective agar plates, as previously described^7^.

### Outgrowth and selection for homology-driven recombination mutants

Following GBS transformation with an editing plasmid, a single colony was resuspended in 20 μL 1x phosphate buffered saline (PBS). 2 μL of this resuspension was used as template for a colony PCR reaction to confirm presence of the intended plasmid. Simultaneously, the remaining 18 μL of the PBS-resuspended sample was seeded into a 10 mL TS+erythromycin 5 μg/mL culture, which was allowed to grow stationary at 37 °C for 6-8 hours.

Following this 6-8-hour outgrowth, 1:10 and 1:100 dilutions of the culture were prepared (dilute in TS+erythromycin). 150 μL of the following cultures were then plated on separate TS+erythromycin 5 μg/mL agar plates: undiluted, 1:10, and 1:100, undiluted+aTC, 1:10+aTC, and 1:100+aTC. For the aTC-containing cultures for plating, we add 6 μL of a 2 mg/mL aTC stock to the 150 μL of sample being plated (for ease, the aTC stock can be added directly into the pooled plating sample on the agar, then the mixture sterilely spread across the surface).

These plates are then allowed to grow for 36 hours at 37 °C. A multiple-log fold difference between colony counts on the aTC and non-aTC plates after growth is indicative of successful Cas12a selection against wild-type GBS. At least 8 colonies are selected and each resuspended in 20 μL 1x PBS. 2 μL of this resuspension is used in a colony PCR reaction to access genotype. Simultaneously, the remaining 18 μL of the PBS-resuspended sample was plated for single colonies on a TS+erythromycin 5 μg/mL plate. Colonies with desired genotype by colony PCR are chosen. Using an isolated colony from the plate struck for single colonies, resuspend a colony in 20 μL 1x PBS and use 2 μL of this resuspension in a second confirmatory colony PCR of genotype. Once confirmed, purify PCR product from gel for sanger sequencing and seed remain 18 μL of colony resuspension into a 40 mL TS broth culture to begin plasmid curing.

### Outgrowth and selection for free end joining mutants

To use pGBSedit, without an editing cassette, to generate free end joining mutants, the same experimental procedures are followed as with generation of a homology-driven recombination mutant. However, because free end joining mutants can be the result of indels that are too small for easy detection by comparing PCR bands on an electrophoresis gel, we first perform a colony PCR screen on approximately 16 colonies, checking for the presence or absence of the wild type Cas12a target (the sequence complementary to the pGBSedit protospacer). We use the target site as the forward PCR primer and choose a reverse primer approximately 500 kB downstream. From this screen, we select candidate colonies that do not generate bands. These are the candidate mutant strains. Then we perform a second colony PCR reaction on the candidate mutant strains, using forward and reverse primers approximately 500 kB from the target site in each direction. Colonies that yield bands on this second reaction can be genotyped using Sanger sequencing of the amplicon or whole genome sequencing to identify and characterize free end joining mutations..

### CFU quantification of Cas12 selective force (Fig 3B)

14 individual TS+5 μg/mL erythromycin cultures were inoculated using GBS harboring the pGBSedit CovR Deletion plasmid and grown for 8 hours. Serial dilutions and CFU plating on TS+5 μg/mL erythromycin plates for each overnight culture were performed to access total viable CFU/mL. Simultaneously, 100 μL of each overnight culture was plated onto TS+5 μg/mL erythromycin+500 ng/mL aTC at both 1:10 or 1:100 dilutions. After 36 hours growth, CFU counts on aTC plates were taken and compared to total number of viable CFU plated from the starting culture

### Measurement of Mutagenesis frequency (Fig 3C)

pGBSedit CovR Deletion plasmid was transformed into electrocompetent GBS in 6 biological replicates. 1 colony from each transformation was grown and aTC selected per previously reported method. 16 colonies from each of the 6 biological replicates were selected and screened for genotypes by NEB OneTaq colony PCR (Cat. M0482L). Genotypes were counted based by 1% agarose gel electrophoresis PCR band sizes (3048 bp WT and 2358 bp covR KO).

### Colony PCR

Colony PCR reactions were performed using NEB OneTaq Master Mix (Cat. M0482L) and custom-prepared ssDNA oligonucleotide primers. Wild type isogenic GBS colonies were used as positive control samples; sterile water was used as template in negative control samples. All templates for reactions are colonies that were first resuspended in 1xPBS. Manufacturer instructions were followed, including proper calculation of primer annealing temperatures and PCR extension time.

### Outgrowth and induction of knockdown strains

GBS harboring the pGBScrispri plasmid are grown in TS broth supplemented with 5 μg/mL erythromycin to maintenance the plasmid. For phenotypic assays, induction of knockdown is achieved by the addition of 250 ng/mL anhydrotetracycline (stock 2 mg/mL in water) at inoculation. Cultures inoculated from isolated single colonies. Cultures are grown stationary at 37°C for 18 hours.

### pGBSedit plasmid curing from confirmed mutants

40 mL TS broth cultures were inoculated using an isolated colony that has been genotypically confirmed and grown stationary at 37°C overnight. A second passage in 40 mL of TS broth is performed using 100 μL of previous culture as inoculum. 1:10,000 and 1:100,000 dilutions of the grown second passage were plated on TS agar plates. 8 colonies selected were dual patched onto TS and TS+5 μg/mL Erm agar plates to identify antibiotic sensitivity.

### Fluorescence assay

10 mL TS broth cultures were inoculated using an isolated colony and grown overnight at 37°C shaking. Shaking cultures were used to ensure oxygen is present during maturation of GFP chromophore formation. Overnight cultures were spun at 4000rpm for 5 minutes and washed twice with PBS before being OD600 normalized to 1 in PBS. 10 mL of OD600 normalized cultures were spun down at 4000rpm for 5 minutes and resuspended in 1 mL of PBS to 10x concentrate. 200 μL of each sample was loaded in technical triplicate into a black optical bottom 96-well plate. GFP was excited at 488 nm and emission was observed at 515 nm with a 515 nm filter.

### RNA purification and qRT-PCR

10 mL TS+5 μg/mL erythromycin broth cultures were inoculated using an isolated colony and grown to mid-log (OD600 of 0.6) before being split into two 5 mL cultures; not treated and 250 ng/mL anhydrotetracycline supplemented.

Cultures were incubated at 37°C for 1 hour post treatment before being spun down at 4000rpm for 5 minutes and resuspended in 1 mL of RNAprotect Bacterial Reagent (Qiagen Catalogue: 76506). Bacteria were incubated in RNAprotect at room temperature for 5 minutes before being pelleted in microcentrifuge tubes at 15,000rpm for 2 minutes. Excess RNAprotect was removed and pellets were frozen and held at -80°C until RNA extraction.

RNA protected pellets were thawed on ice and RNA was extracted using the Qiagen RNeasy Kit (Cat. 74104) per manufacturer protocol. RNA was eluted with warmed (95°C)ultra-pure water in place of elution buffer. DNA was removed with Invitrogen DNA-free Kit (Cat. AM1906) per manufacturer protocol. cDNA was made with AppliedBiosytems High-Capacity cDNA Reverse Transcription Kit (Cat. 4374966) per manufacturer protocol. cDNA and reverse transcription negative were 1:10 diluted with ultra-pure water before 1 μL was used as template in 10 μL qPCR reactions. Bio Rad SsoAdvanced Universal SYBR Green Supermix (Cat. 1725272) was used for all qPCR reactions.

### Hemolysis assays

10 mL of human whole blood was donated, heparinized and spun down for 10 minutes at 2000 rpm. Serum was removed with a serological pipette and 5 mL of Hanks buffers saline solution (HBSS) is added to the packed red blood cells via inversion. Packed red blood cells (RBC) were spun for 10 minutes at 2000rpm and the supernatant was removed with a serological pipette. 500uL of packed RBCs were added to 50 mL of HBSS to achieve a 1% packed RBC solution. 10 mL cultures of GBS harboring the pGBScrispri plasmid were grown in TS broth supplemented with 5 μg/mL erythromycin and either with or without 250 ng/mL of anhydrotetracycline. Cultures were grown stationary at 37°C for 18 hours. Cultures were pelleted at 4000rpm and resuspended in 10 mL of PBS to wash. 10 mL PBS resuspensions were OD600 normalized to 1 using a cuvette. 10 mL of OD normalized A909 cultures were concentrated 10x to an OD600 of 10 by pelleting and resuspending in 1 mL of PBS. 10/84 cultures were 1:10 diluted in PBS after OD normalization. Hemolysis assays were performed with 100 μL of 1% packed RBCs and 100 μL of bacterial suspension incubated at 37°C for either 15 minutes (10/84) or 1 hour (A909). Hemolysis assays were performed in 96 well V-bottom plates and in technical triplicate.

After hemolysis, V-bottom plates were spun down at 2000rpm for 2 minutes and supernatants (100 μL) were collected for each sample and absorbance at 415 nm was used to measure released hemoglobin in the supernatant. Percent hemolysis is calculated relative to a 1% triton control.

### Whole-genome sequencing, scaffolding, and analysis

Genomic DNA samples were prepared for whole-genome sequencing (WGS) using the Illumina DNA Prep tagmentation kit with IDT for Illumina Unique Dual Indexes. Sequencing was performed on an Illumina NextSeq2000 platform with a 300-cycle flow cell to generate paired-end reads (2×150 bp).

A 1-2% PhiX control library was included to ensure optimal base-calling accuracy. Initial processing—including demultiplexing, read trimming, and analytics—was performed using Illumina DRAGEN software (version 3.10.12)^48^. Quality control included FastQC metrics evaluation.

Further processing and genomic assembly utilized a custom bioinformatics pipeline. Reads underwent stringent quality filtering, trimming, and duplicate grouping using BBTools. Filtered reads were aligned to the A909 GBS strain reference genome to create a high-confidence consensus sequence, requiring a minimum depth of 10 reads at consensus sites.

Reads were assembled into contigs using SPAdes^49^ in careful mode. Contigs were scaffolded onto the consensus reference using RagTag^50^ , followed by patching to resolve scaffold gaps and inconsistencies. The primary scaffold from each assembly was extracted and annotated using the Prokaryotic Genome Annotation Pipeline (PGAP)^51^. Pipeline scripts and dependencies were managed within a custom conda environment and run locally.

## Supporting information

Supplemental protocols

## Acknowledgements

This work was supported by the National Institutes of Health (NIH) R21AI178067, R01AI182835, and R01AI177991 (T.A.H.), a UPMC Children’s Hospital of Pittsburgh Research Advisory Council graduate student grant (G.H.H.), and Centers of Biomedical Research Excellence (CoBRE) Grant P20GM125504 (J.C.). Figure panels throughout were generated using BioRender.

## Supplemental materials

**Supplemental Figure S1:**
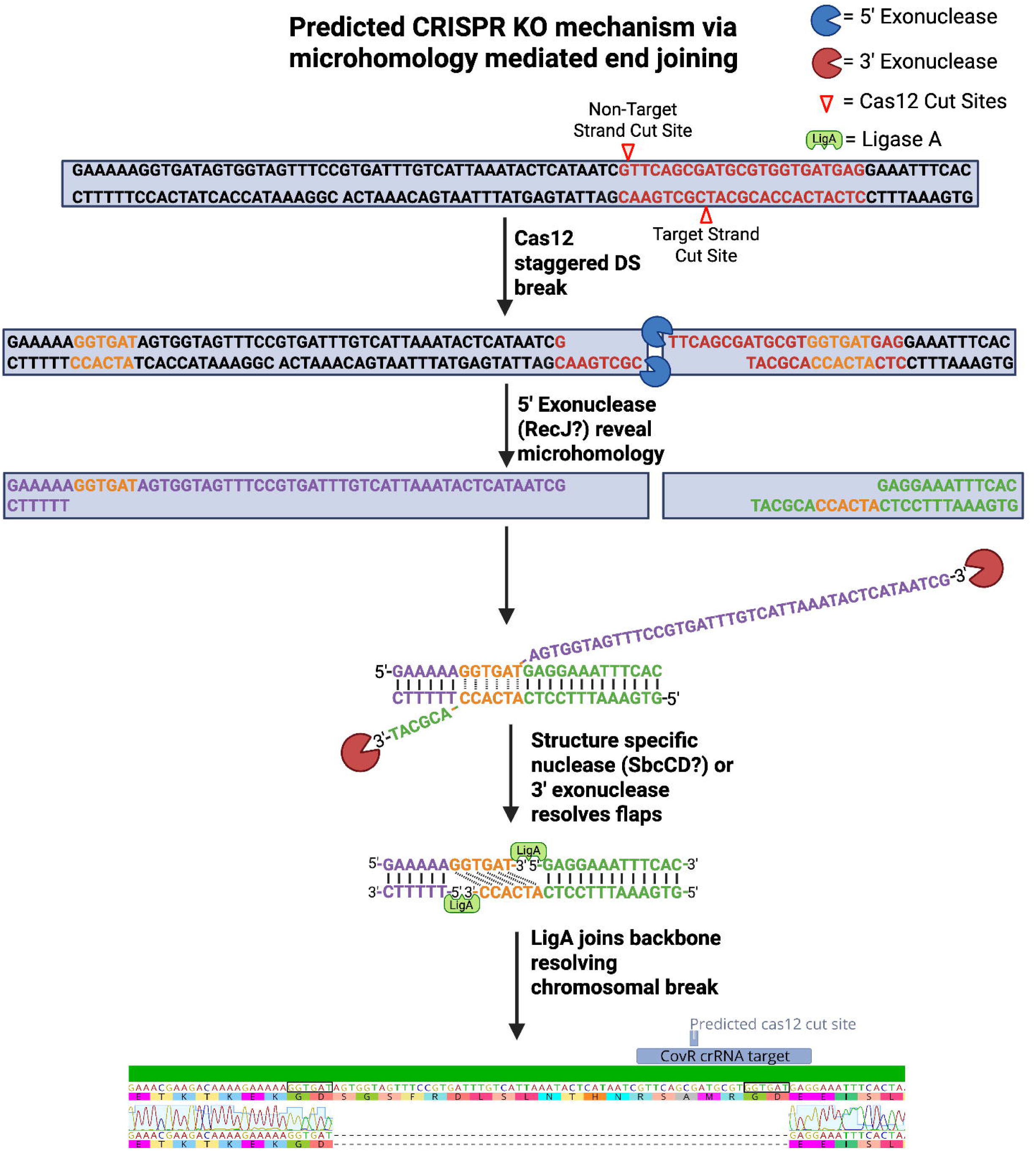
Proposed mechanism of alternative end-joining in GBS.

**Supplemental Extended Protocol**: Detailed instructions for using pGBSedit and pGBScrispri.

## Public resources availability

The following plasmids for GBS Cas12a editing, CRISPRi, and complementation are available at Addgene.

| Addgene Plasmid Number | Plasmid Name |
| --- | --- |
| 223200 | pGBSedit Sham |
| 223201 | pGBScrispri Sham |
| 223202 | pGBScomp High |
| 223203 | pGBScomp<br>Low |

