## Supplemental protocols for "A Cas12a Toolbox for Rapid and Flexible Group B *Streptococcus* Genomic Editing and CRISPRi"

### Supplemental Protocol

### GBS Cas12a Genome Editing Protocol with pGBSedit

**Visual Aids:** *See end of document*

**Visual aid 1.** pGBSedit/CRISPRi modular crRNA cloning

**Visual aid 2.** pGBSedit cloning in homology arms

**Visual aid 3.** pGBSedit expedited/minimal cloning approach using a gene fragment

**Materials needed:**

- Primers: Insert Screen Fwd, Insert Screen Rev, pGBSedit crRNA Screen Rev
  - Insert Screen Fwd: AAAAAGATGCCAGTGTGCTG
  - Insert Screen Rev: CCAGTGCCAAGCTTGCATGT
  - pGBSedit crRNA Screen Rev: GCTATCTTCGTCATAGTTACCT
- Gibson Assembly primers needed if using Gene Fragments
  - GeneFrag Insert Fwd: GCAATAAAACTTATTCGGATCCGTCGTTTTACAACGTC
  - GeneFrag Insert Rev: AGCTGACGGTCAGAGAGAAAGGATTTCTCACATAAAATAGAGG

- Restriction enzymes: *XhoI*, *Esp3I*, *DpnI*
- Gibson assembly master mix
- T4 DNA Ligase and T4 DNA ligase buffer
- T4 Polynucleotide Kinase
- aTC (anhydrotetracycline)
- PCR and Gel purification kits (Qiagen)
- Chemically competent *E. coli* and electrocompetent GBS

**General steps:**

- Design and insert new crRNA against protospacer in target gene into pGBSedit Sham
- Design allelic exchange homology arms to insert into pGBSedit with new crRNA via Gibson Assembly
- Design and order primers to screen chromosomal locus for WT vs KO strains
- Gibson Assemble homology arms into pGBSedit plasmid with new crRNA and transform into *E. coli*
- Patch and confirm patches with Insert via Insert Screen Fwd and Insert Screen Rev primers
- Transformation into GBS (*requires electrocompetent GBS*)
- Grow GBS strain with pGBSedit + editing cassette, adding cas12a inducer to select for chromosomal edit
- Colony PCR patches with chromosomal screening primers to identify mutants
- Serial passage without antibiotic to cure genome editing plasmid
- Sequence mutant and stock to strain list
- Optional: Complement KO with pGBScomp

**Notes:**

- Tryptic Soy media is used through this protocol, but any nutrient rich media (THB, THY, BHI) is sufficient.
- A foundational understanding of allelic exchange homology directed repair based genomic editing and access to Geneious Prime/Genome viewer are assumed in this protocol.
- Anhydrotetracycline (inducer of the Cas12a Pxyl/tet promoter) is light-sensitive and any step involving it should be kept away from the light

### Designing pGBSedit Gene Knockout Plasmid

1. **Select a gene to edit in Geneious Prime**
   a. Confirm this gene is not essential

- **Design Cas12a crRNA against gene for pGBSedit**
  a) Copy gene to knockout into CRISPick to find protospacer (https://portals.broadinstitute.org/gppx/crispick/public)
  b) Ref Genome does not matter
  c) Mechanism is CRISPR KO
  **d) <u>Enzyme is AsCas12a (TTTV)</u>**
  e) Keep Quota at 5
  f) Check box for "Report unpicked sequences"
  g) Save/Copy output file into Excel
  h) Choose sgRNA sequence in the first 20% of the gene
     - The reported sgRNA sequence is the same thing as crRNA sequence
     - Make sure "quota is met" and nothing is wrong with target sequence (no poly TTTT)
  i) Copy sgRNA/crRNA into a new excel page
- **crRNA will be ordered as a Fwd&Rev Oligos from IDT**
  a) Importantly, Oligos will have sticky ends added to match *Esp3I* cut sites on pGBSedit Sham
     - For Fwd Oligo: concatenate "TAGAT" to beginning of fwd sgRNA sequence and "A" to end
     - For Rev Oligo: concatenate "AAATT" to beginning revcom sgRNA sequence and "A" to end

**<u>EXAMPLE OLIGOS</u>: If CRISPick sgRNA sequence chosen is** CCGTCTAACGTTGCATGATAATC

- **RevCom:** GATTATCATGCAACGTTAGACGG
- **Fwd Oligo (TAGAT + sgRNA + A):** TAGATCCGTCTAACGTTGCATGATAATCA
- **Rev Oligo (AAATT + revcom sgRNA + A):** AAATTGATTATCATGCAACGTTAGACGGA

#### <u>Design homology exchange cassette</u>

1. **Select >500 bp of sequence upstream AND downstream of gene of interest to create homology arms**
   a. If your gene has overlapping coding region with another gene, your homology arms must include the overlapping region to not disrupt function of overlapping gene
   b. Larger homology arms should be used if inserting/deleting large sections
2. **Copy each homology arm as a separate fragment into in silico Gibson assembly (NEBuilder) along with pGBSedit (linearized with *XhoI* or via PCR)**
   a. If designing a gene replacement instead of clean knockout, Gibson assemble the gene you want to replace with between the homology arms
   b. You can alternatively design your own Gibson assembly using PCR linearization of pGBSedit if desired
3. **NEBuilder will provide primers required for effective 3-piece Gibson assembly of Upstream homology arm + Downstream homology arm+ linearized pGBSedit**
   a. Be sure to save output PDF for record keeping
   b. Download expected GA product from NEBuilder and examine in Geneious to ensure plasmid is correctly assembled

#### <u>Design Chromosomal Screening Primers</u>

*These primers will allow you to screen and identify mutants.*

1. **Design Forward Primer roughly 750 bp upstream from the start gene you want to knockout and reverse primer roughly 750 bp downstream of the end of the gene.**
    a. It is critical that these primers DO NOT anneal to regions found within the homology arms on the plasmid.
    b. Note that OneTaq annealing temps for these should be above 48 °C . Temps below this have lower efficiency when screening GBS colonies.
2. **Order primers from IDT**

#### Start of Bench work

##### Insert Cas12a new crRNA into pGBSedit: See Visual Aid 1

#### Using Modular Cloning (Not GeneFragment) Start here:

***If using Gene Fragment, skip to after step 4 of "Clone Editing Cassette into pGBSedit"***

1. **Mini-Prep pGBSedit sham plasmid**
2. **Digest pGBSedit sham with *Esp3I***
    **a.** 40 µL reaction digest at 37C for 2 hours
        i. ~3 µg of plasmid
        ii. 4 µL of 10x rCutsmart
        iii. 3 µL of *Esp3I*
        iv. H20 up to 40 µL
3. **Run on gel and gel purify digested plasmid**
    **a.** Qiagen gel purification kit
4. **Rehydrate crRNA Fwd & Rev Oligos**
    a. Normal 100 µM stocks
5. **Phosphorylate crRNA oligos**
    a. 25 µL reactions at 37 °C for 1 hour
        i. 1 µL Fwd oligo stock and 1 µL Rev oligo stock
        ii. 2.5 µL of 10x T4 ligase buffer
        iii. 0.5 µL T4 Polynucleotide kinase
        iv. 20 µL $H_2O$
6. **Anneal phosphorylated crRNA oligos**
    a. Add 2.5 µL of 1 M NaCl to phosphorylation reaction
    b. Heat up to 95 °C and hold for 5 minutes
    c. Allow to cool SLOWLY to 4 °C
        i. I do 30 minutes at room temperature and 30 minutes at 4 °C (refrigerator)
7. **Ligate annealed crRNA with *Esp3I* digested pGBSedit sham**
    a. 20 µL reaction at 16 °C overnight OR at least 2 hours
        i. 5 µL *Esp3I* digested pGBSedit sham
        ii. 2 µL of annealed crRNA oligos
        iii. 2 µL of 10x T4 ligase buffer
        iv. 1 µL T4 Ligase
        v. 9 µL H20
8. **Transform chemically competent DH5a with ligation product**
    a. 5 µL of ligation product
    b. Standard heat shock method, refer to *E. coli* Transformation protocol
    c. Incubate in SOC media for 1 hour and plate 150 µL onto LB+Erm300 µg/mL
9. **Check for colonies the next day**

    a. Patch 4 colonies onto a new LB+Erm300 µg/mL plate for screening

**10. Screen patches for new crRNA**

    a. Using your Fwd crRNA oligo and pGBSedit crRNA Screen Rev
    b. 25 µL OneTaq Colony PCR reactions
        i. 12.5µL of OneTaq 2xMM
        ii. 1 µL of your Fwd crRNA oligo (10 µM)
        iii. 1 µL of pGBSedit crRNA Screen Rev (10 µM)
        iv. 10.5 µL of $H_2O$
        v. Resuspend a very small amount of patches to screen into reactions
    c. Aneal at 47 °C and extend for 30 seconds
    d. Colonies with new crRNA will appear as a ~300 bp band when run on a gel. Colonies with parental (sham) plasmid will NOT yield a band.

**11. Save Ecoli strain pGBSedit with new crRNA to stocks**

#### <u>Clone Editing Cassette into pGBSedit: See Visual Aid 2</u>

1. **Mini-prep pGBSedit with desired targeting crRNA**
2. **Linearize pGBSedit containing new crRNA**
    a. **Option 1: PCR linearize pGBSedit**
        i. 50 µL Q5 PCR with PCR Linearize Fwd and Rev
            **1.** PCR Linearize Fwd: ACATGCAAGCTTGGCACTGG
            **2.** PCR Linearize Rev: CATCTAGAGCTCCGGACGTC
            **3.** Anneal 68 °C, Extension 5 minute 30 seconds
    b. **Option 2: Restriction digest pGBSedit w/ crRNA with *XhoI* for 2 hours at 37 °C**
        i. 40 µL reaction digest at 37C for 2 hours
            **1.** ~3 µg of plasmid
            **2.** 4 µL of 10x rCutsmart
            **3.** 3 µL of *XhoI*
            **4.** $H_2O$ up to 40 µL
3. **PCR Purify (or gel purify) the linearized plasmid**
    a. PCR purification can be used in this step since *XhoI* only makes a single cut (unlike *Esp3I*)
4. **Prepare homology arms for Gibson Assembly into linearized pGBSedit w/ crRNA**
    a. **<u>IF homology arms are generated via PCR (for 3 piece Gibson assembly)</u>**
        i. Q5 PCR homology arms off WT GBS gDNA
        ii. Use NEBuilder recommended annealing temp
        iii. Run PCR products on gel and purify PCR bands

#### <u>Using Ordered Gene Fragment Start Cloning Here: See Visual Aid 3</u>

##### <u>*Steps with red bullets only necessary when using a gene fragment*</u>

- Use the GeneFrag Insert Fwd&Rev primers to PCR linearize the plasmid
    - 50 µL reaction, $T_m$=59 °C , Ext: 6 min
        - 25 µL Q5 2xMM
        - 20 µL $H_2O$
        - 2 µL Fwd Primer (10 µM)
        - 2 µL Rev Primer (10 µM)
        - 1 µL 1:100 diluted pGBSedit Sham
- *DpnI* digest PCR product
    - 60 µL reaction, 37 °C, 1 hour
        - 50 µL of finished PCR product

        - 6 µL of rCutsmart
        - 2 µL of $H_2O$
        - 2 µL of *DpnI*
- PCR Purify (or gel purify) the *DpnI* digested PCR
    - I rehydrate in only 30 µL $H_2O$ to help concentrate the DNA
- Rehydrate Gene fragment
    - Add 20 µL of $H_2O$
    - Warm shake at 37 °C for 1 hour (or vortex and spin down every ~10 minutes while heating)
- Proceed to Step 5 of "Clone Editing Cassette into pGBSedit"
    - Gibson assembly with PCR linearized pGBSedit and rehydrate gene fragment

5. **Set up 20 µL Gibson assembly reaction**
    a. 10 µL of 2xHiFi cloning MM & Equal mix (by volume) of all fragments up to 20 µL.
    b. Incubate at 50 °C for 1 hour (2 hours if more than 3 fragments)
6. **Transform 5 µL of Gibson assembly reaction into chemically competent Dh5a *E. coli***
    a. Standard heat shock method, refer to *E. coli* Transformation protocol
    b. Incubate in 500 µL of SOC media for 1 hour and plate 150 µL onto LB+Erm300
7. **Screen *E. coli* colonies for pGBSedit +homologous exchange cassette with colony PCR**
    a. Use Insert Screen Fwd and Insert Screen Rev primers
    b. 25 µL OneTaq Colony PCR reactions, Annealing temp 54 °C, 2-minute extension
        i. 12.5 µL of OneTaq 2xMM
        ii. 1 µL Insert Screen Fwd (10 µM) & 1 µL Insert Screen Rev (10 µM)
        iii. 10.5 µL of $H_2O$
        iv. Resuspend a very small amount of patches to screen into reactions
            1. Best to resuspend patch into small volume of PBS and use 1 µL as template in this reaction
    c. Alternatively, mini-prep and send for whole plasmid sequencing
8. **Save *E. coli* strain pGBSedit w/ editing cassette**

#### GBS Genomic Editing

1. **Mini-prep pGBSedit w/ editing cassette**
    a. Need a good concentration >100 ng/µL for decent efficiency GBS transformation
    b. I will always whole plasmid sequence before using the plasmid
2. **Transform electrocompetent GBS with mini-prep**
    a. Use GBS electroporation protocol
    b. Use 5 µL of plasmid mini-prep in transformation
    c. Plate entire recovery across 3 TS+Erm5 plates
3. **<u>In the morning of the next day</u>, identify transformant colonies**
    a. Start three 5 mL TS+Erm5 cultures using colonies of transformation plate and grow at 37 °C for 6-8 hours
        i. ***I do NOT do overnight growths at this step. We have seen extended exposure/multiple passages with the chromosomal targeting pGBSedit result in a higher rate of aTC resistant colonies that are NOT the desired mutant.***
4. **After 6-8 hours of outgrowth, plate onto TS+Erm5+500 ng/mL aTC**
    a. Plate 100 µL of bacterial cultures at full concentration and 1:10 dilution, onto TS+Erm+aTC and TS+Erm plates
        i. *Often after 6-8 hours of growth there is still no visible growth in the culture. This is ok. We see best results come from plating at very early growth.*

   ii. *The difference in colony number between aTC and no aTC will give you an idea of how well your selection worked. If STRONG selection is not observed on aTC plates, restart at GBS transformation step!*

   b. To add aTC to the surface of a plate, add 6 µL of aTC stock (*2 mg/mL)* to your bacteria being plated before spreading with beads.
      i. *500 ng/mL concentration is calculated by spreading 6 µL of stock onto surface of 25 mL agar plates*
      ii. *We have had aTC stability issues when added directly to molten agar*
   c. Allow to grow at 37 °C **protected from light for 36 hours**
      i. I keep plates under a box in the incubator
      ii. The extra 24 hours of growth helps visual differentiation of healthy colonies

5. **Isolate single colonies with healthy phenotypes and resuspend in 20 µL of PBS and plate**
   a. Identify colonies with healthy phenotypes
      i. *Avoid very small colonies or ones with "ghostly"/transparent or other unhealthy phenotypes*
      ii. *Do not avoid orange colonies, this occurs because aTC exposure modestly induces cyl expression*
   b. Select 8 colonies and resuspend each colony in 20 µL of PBS.
      i. *It is imperative that you isolate a single colony off the plate at this step.*
   c. Plate 15 µL onto a fresh TS+Erm plate, streak for single colonies
      i. You will need single isolated colonies for future steps
6. **Colony PCR colonies with your genes designed chromosomal screening primers**
   a. Colony PCR colonies using 2 µL of resuspension from step 5.
   b. Be sure to include a WT control
   c. 25 µL reactions. Use NEB TM calculator for OneTaq to determine annealing temp
7. **Run Colony PCR reactions on a 1% agarose gel**
8. **Identify genotypes of patches by size of PCR product bands**
   a. *Remember the chromosomal screening primers are designed to create a different sized product (smaller for knockouts) bands for patches with edited genomes when compared to those that are WT (unedited)*
      i. *I often only see ~10-50% colonies having edited genotype depending on the gene.*
      ii. *Many of your bands will appear WT, these are colonies that mutated the crRNA target site.*
      iii. *If you see both the WT and mutant band, this is likely because of mixed populations at step 5*
   b. *If you see very high rates (>25%) of failed PCR, try designing new primers.*
      i. *This happens regularly with GBS colony PCR primers*
   c. *If ALL patches appear WT return to step step 5 and screen more colonies. Some genes with high fitness costs may be very rare and impossible to find without an extra selective marker (such as an added ABX resistance gene)*
   d. *If you have screened more than 24 colonies, return to the GBS transformation step (Step 2)*
      i. If efficiency is still too low to find your mutant here are a few ways to boost it/common issues:
         1. Increase size of homology arms to 1 kbp each
         2. Choose new crRNA with higher GC content (should be at least >30%)
         3. Plasmid sequence your complete pGBSedit and ensure no errors
9. **Once the PCR identified a mutant genotype, return to the plate for that strain from step 5**
   a. Resuspend an isolated single colony from this plate into 20 µL of PBS
   b. Use 2 µL in a second colony PCR set up exactly the same as in step 6
      i. *Goal is to confirm that colony you will be proceeding with has the desired genotype*
      ii. *Step 9 was added to help reduce the seemingly incidence of WT reversion caused by mixed populations*
   c. Use other 18 µL to seed 40 mL TS culture for plasmid curing

### pGBSedit plasmid curing from confirmed knockout

1. **Start a 40 mL culture of TS with a PCR confirmed knockout strain.**
    a. This is the same culture from the previous step (see step 9c)
    b. *Goal is to get GBS to cure the pGBSedit plasmid so do not add Erm!*
    c. Let grow stationary at 37 °C either 8 hours or overnight
2. ***Start a second passage in TS***
    a. 40 mL culture
    b. Inoculate with 100 µL of the previous culture
    c. Let grow away overnight stationary at 37 °C
3. **Plate the following dilutions of the overnight culture onto a TS plate**
    a. 1:10,000 and 1:100,000
    b. Allow to grow overnight
4. **Dual patch single colonies onto TS and TS+Erm5 plates**
    a. Be sure to patch colonies onto the TS plate first and TS+Erm5 plate second
    b. 8 patches are usually more than sufficient
    c. Grow plates overnight
5. **Identify a patch that is Erm sensitive**
    a. This strain has cured its plasmid and is no longer resistant to the antibiotic
6. **Final confirmation of knockout strain**
    a. Screen plasmid cured knockouts with colony PCR
    b. Gel extract and sequence these genotype PCRs
        i. Alternatively, gDNA extract and WGS
7. **Save the plasmid cured knockout strain to stocks**
    a. Strain should be WGS before use

### Complementing KO strains with pGBScomp

**Materials needed:**

- Restriction enzymes: *SalI*
- *E. coli* with pGBScomp-High (available on Addgene Catalog: 223202)
- PCR and Gel purification kits (Qiagen)
- Gibson assembly master mix
- Chemically competent *E. coli* and electrocompetent GBS
- Primers to PCR gene to complement

#### Designing pGBScomp Gene Complement Plasmid

1. Select gene you want to complement and acquire sequence.

CovR Example:
ATGGGTAAAAAGATCTTAATAATCGAAGATGAGAAAAATTTAGCTCGCTTCGTCTCGTTAGAACTACTACATGAAGGATATGATGTTGTTG
TTGAAACAAACGGTCGTGAAGGATTGGACACAGCATTAGAAAAAGATTTTGATTTGATTCTACTGGATTTAATGCTTCCAGAGATGGATG
GTTTCGAAATCACACGTCGCCTGCAGGCTGAAAAAACAACCTATATCATGATGATGACAGCACGTGATTCTGTTATGGATATTGTAGCTGG
TCTTGATCGTGGAGCAGATGATTATATTGTTAAGCCGTTTGCAATCGAAGAATTATTAGCACGTGTTAGAGCGATTTTCCGACGCCAAGA
AATTGAAACGAAGACAAAAGAAAAAGGTGATAGTGGTAGTTTCCGTGATTTGTCATTAAATACTCATAATCGTTCAGCGATGCGTGGTGA
TGAGGAAATTTCACTAACAAAACGTGAATTTGATTTGTTGAATGTCTTGATGACAAATATGAATCGTGTTATGACACGAGAAGAGTTGCT
AGAACATGTTTGGAAATACGATGTGGCAGCAGAGACAAACGTTGTTGATGTTTATATCCGTTACCTAAGAGGTAAAATTGATATCCCAGG
TCGTGAATCATATATTCAAACTGTTCGCGGAATGGGCTATGTGATTCGTGAAAAATAA

2. Design PCR primers to PCR gene

CovR Example:

CovR Fwd: ATGGGTAAAAAGATCTTAATAATC

CovR Rev: TTATTTTTCACGAATCACATAG

3. Add pGBScomp-High matching overlaps to primers

CovR Example:

pGBScomp-CovR Fwd: **tgacaatgatgttggatccg**ATGGGTAAAAAGATCTTAATAATC

pGBScomp-CovR Rev: **tcgataagcttggctgcagg**TTATTTTTCACGAATCACATAG

4. Order primers with overlaps as oligos from IDT

#### Cloning pGBScomp

1. Grow *E. coli* strain with pGBScomp-High plasmid
    a. 5mL culture of LB + Erm300 µg/mL
    b. Shaking at 37 °C
2. Mini-prep pGBScomp-High Plasmid

3. *SalI* Digest pGBScomp-High
    a. 40 µL reaction digest at 37 °C for 2 hours
        i. ~3 µg of plasmid
        ii. 4 µL of 10x rCutsmart
        iii. 3 µL of *SalI*
        iv. $H_2O$ up to 40 µL
4. PCR Purify (or gel purify) the digested plasmid
5. PCR gene of interest using your designed Gibson assembly primers
    - 50 µL reaction
        - 25 µL Q5 2xMM
        - 20 µL L $H_2O$
        - 2 µL Fwd Primer (10 µM)
        - 2 µL Rev Primer (10 µM)
        - 1 µL L Template (usually diluted gDNA)

6. Gel purify PCR product
    a. I use 30 µL of water for the final dilution to get a higher final concentration
7. Set up 20 µL Gibson assembly reaction
    a. 10 µL of 2xHiFi cloning MM, 5 µL of *SalI* digested pGBScomp-High, 5 µL of gene of interest PCR
    b. Incubate at 50 °C for 1 hour
8. Transform 5 µL of Gibson assembly reaction into chemically competent DH5a *E. coli*
    a. Standard heat shock method, refer to *E. coli* Transformation protocol
    b. Incubate in 500 µL of SOC media for 1 hour and plate 150 µL onto LB+Erm300
9. Patch 2 colonies from *E. coli* transformation plates and grow overnight
10. Start a 5 mL LB+Erm300 culture for each *E. coli* patch and grow 37 °C shaking
11. Mini-prep and send plasmid for sequencing
    a. We like Plasmidsaurus
12. Once plasmid sequence in confirmed, transform into electrocompetent GBS
13. Confirm overexpression of complement gene using qPCR or phenotypic assays

#### pGBSedit/CRISPRi modular crRNA cloning

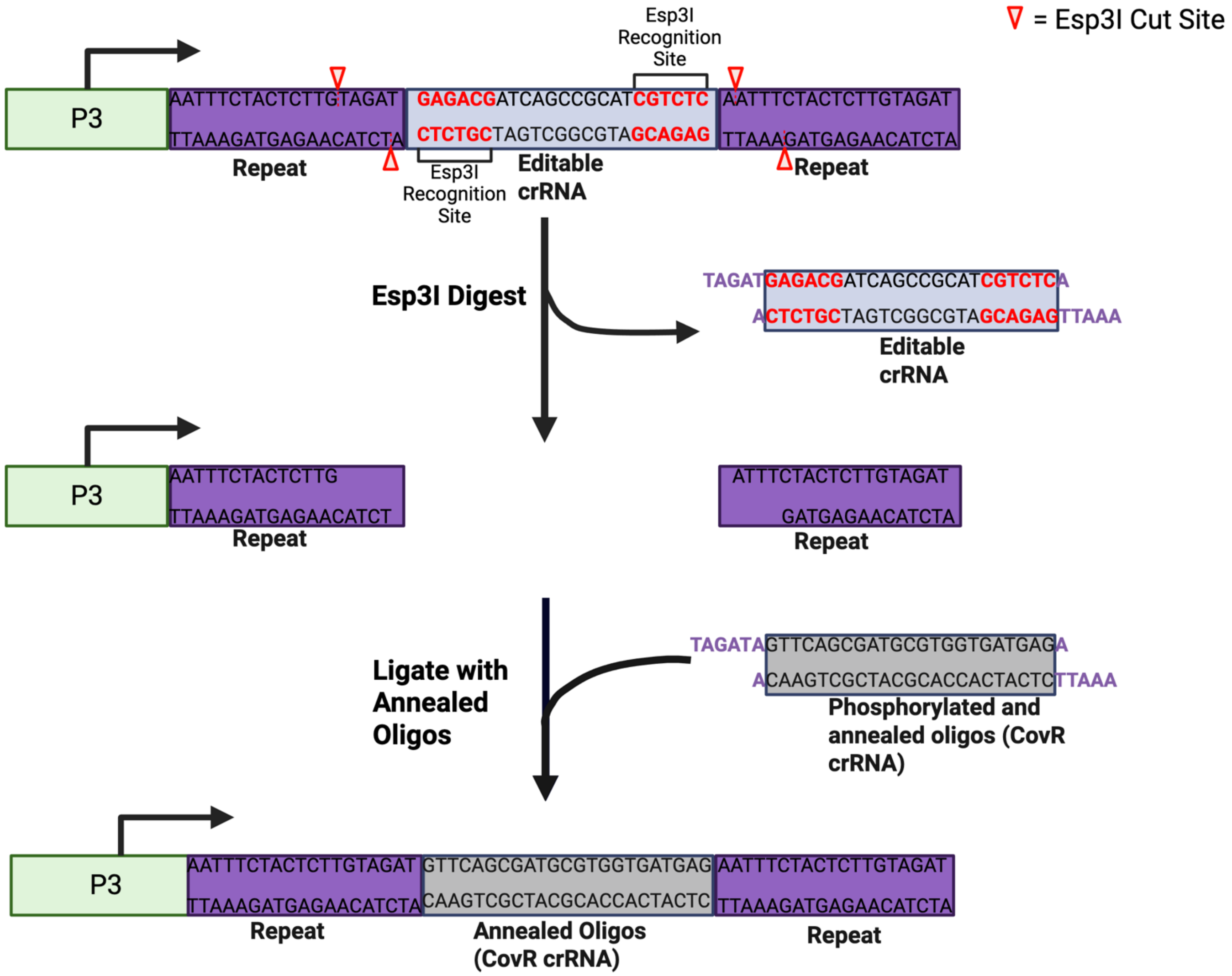


**Visual aid 1:** The pGBSedit plasmid contains a sham crRNA flanked by two *Esp3I* recognition sites. Digestion with *Esp3I* excises the sham crRNA, generating sticky ends suitable for ligation. Custom crRNA spacers can be introduced by annealing and phosphorylating complementary oligonucleotides with matching overhangs and ligating them into the digested vector. This allows rapid and modular replacement of the crRNA for targeting desired sequences, such as with CovR shown here.

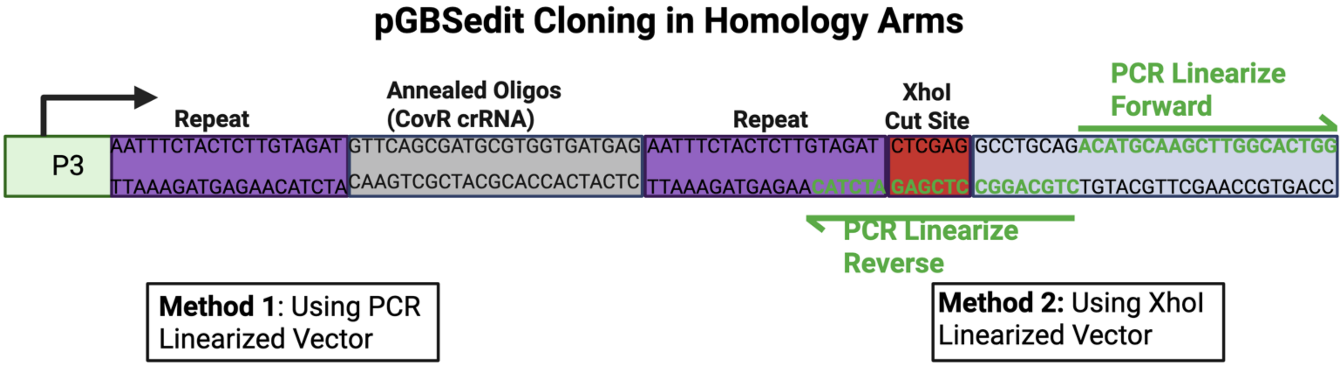


**Visual aid 2:** Two approaches are available for preparing the pGBSedit plasmid for insertion of homology arms via Gibson assembly. Method 1 employs PCR amplification using the indicated forward and reverse primers (green) to generate a linear vector. Method 2 uses restriction digestion at the unique *XhoI* site to linearize the plasmid. Both methods enable a user designed Gibson Assembly for insertion of flanking regions for targeted genome editing.

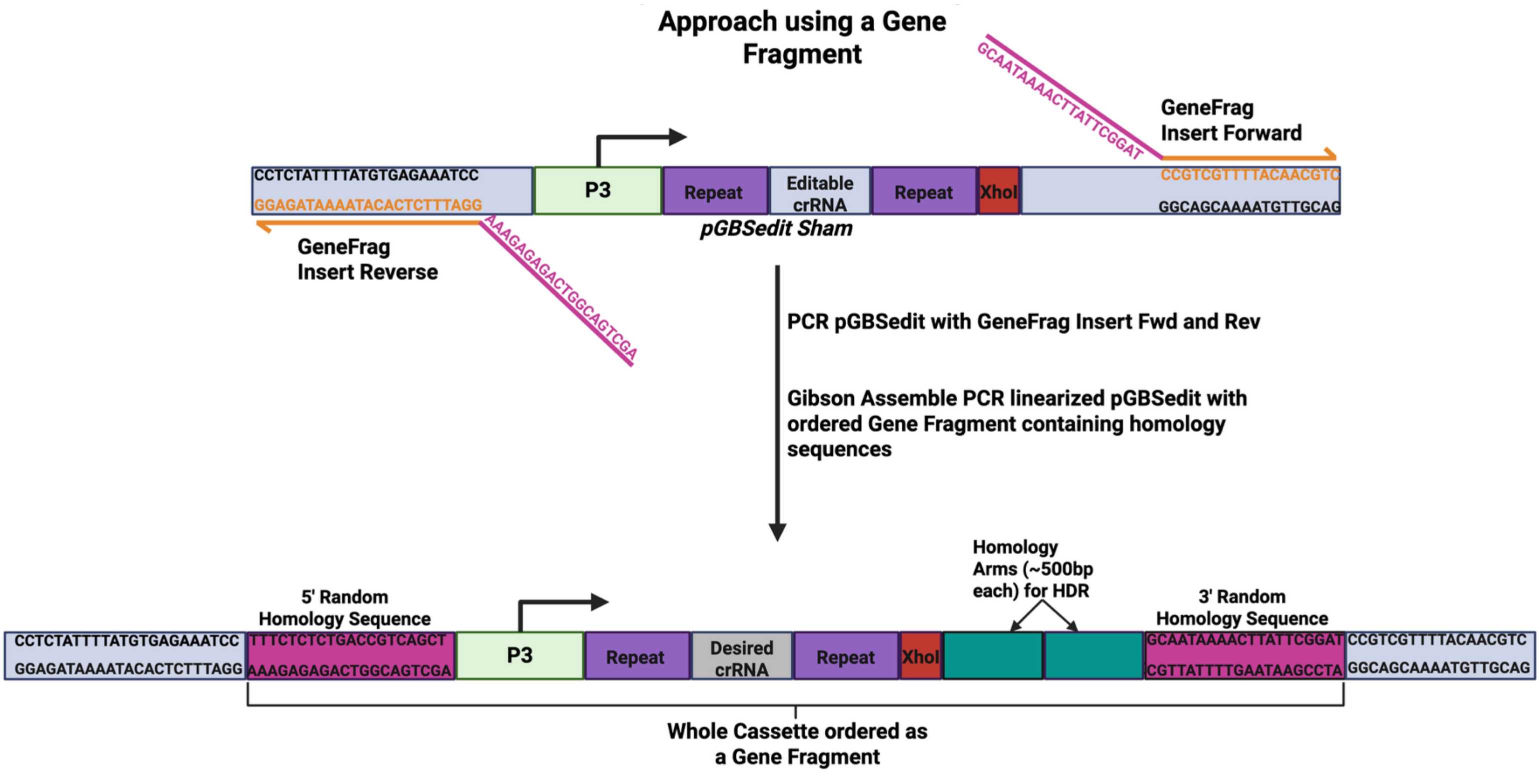


**Visual aid 3:** This expidited/minimal cloning approach enables rapid generation of a complete pGBSedit editing plasmid by bypassing the separate cloning steps for addition of crRNA and homology arms. A single synthetic gene fragment encoding the entire desited editing cassette is ordered. The pGBSedit sham backbone is PCR-linearized using primers (GeneFrag Insert Fwd/Rev) that introduce random homology overhangs (shown in pink). These overhangs allow seamless Gibson Assembly with any gene fragment designed to carry matching flanking sequences, enabling modular integration of any custom editing cassettes.
